# *Culex saltanensis* and *Culex interfor* (Diptera: Culicidae) are susceptible and competent to transmit St. Louis encephalitis virus (Flavivirus: Flaviviridae) in central Argentina

**DOI:** 10.1101/722579

**Authors:** Mauricio D. Beranek, Agustín I. Quaglia, Giovana C. Peralta, Fernando S. Flores, Marina Stein, Luis A. Diaz, Walter R. Almirón, Marta S. Contigiani

## Abstract

Infectious diseases caused by mosquito-borne viruses constitute health and economic problems worldwide. St. Louis encephalitis virus (SLEV) is endemic and autochthonous in the American continent. *Culex pipiens quinquefasciatus* is the primary urban vector of SLEV; however, *Culex interfor* and *Culex saltanensis* have also been found naturally infected with the virus, suggesting their potential role as vectors.

**OBJECTIVE:** The aim of this study was to determine the vector competence of *Cx. interfor* and *Cx. saltanensis* for SLEV from central Argentina in comparison to *Cx. p. quinquefasciatus*.

**METHODS:** Adult female mosquitoes of the three *Culex* species were orally infected by feeding on viremic chicks that had been inoculated with SLEV. Then, abdomens, legs and saliva blood-fed mosquitoes were analyzed by viral plaque assay and the presence of cytopathic effect on the cell culture monolayer.

**RESULTS:** Mosquitoes were permissive to orally acquired infections, to virus dissemination, and transmission of SLEV in the saliva. *Cx. saltanensis* and *Cx. interfor* are potential vectors of SLEV.

**CONCLUSIONS:** Our results demonstrate that in Argentina both *Cx. saltanensis* and *Cx. interfor* are susceptible to SLEV and competent for its transmission. Moreover they are abundant during SLEV epidemic period in urban area, positive for this virus in nature, and found to feed on natural hosts.

This article has been peer-reviewed and recommended by *Peer Community in Entomology*

## Introduction

Infectious diseases caused by vector-borne pathogens constitute health and economic problems worldwide [1]. Mosquitoes are an important group of arthropod vectors. Due to the hematophagous habit of females, many mosquito species are vectors of infectious agents, including viruses (arthropod-borne viruses; ‘arboviruses’) *2+. Arbovirus is maintained by biologic transmission among vectors and hosts. Sometimes this biological transmission is specific and includes few vector and host species such as Chikungunya (CHIKV), dengue (DENV), urban yellow fever (YFV) and Zika viruses (ZIKV). However, most of the arboviruses are generalist and they use many vectors and hosts species such as St. Louis encephalitis virus (SLEV) and West Nile virus (WNV) [3]. The emergence and reemergence of diseases caused by arbovirus are a global phenomenon, in particular, those caused by CHIKV, DENV, WNV and ZIKV [1].

SLEV is an endemic neurothropic flavivirus in temperate and subtropical areas of the New World. This virus is maintained between multiple avian hosts and *Culex* vectors, although incidental infections are possible in humans and other mammals, which are typically dead-end hosts [4]. In 2002, SLEV reemerged in the central area of Argentina and southern Brazil causing neurological diseases in humans. In 2005, the first outbreak occurred in Córdoba City with 47 confirmed cases and nine fatalities. After the 2005 outbreak, additional SLEV outbreaks in Argentina occurred in Parana (2006), Buenos Aires (2010), and San Juan (2011) [5]. Factors that promoted this emergence in Argentina include the introduction of a more virulent SLEV strain into a highly susceptible avian hosts community along with possible land use changes (urbanization, agriculture) [5]. Phylogenetic analyses indicate that the emerging SLEV in US (2015) is related to the epidemic strains isolated during a human encephalitis outbreak in Argentina (2005), suggesting introduction from South America [5].

SLEV, as similar to other arboviruses transmitted by mosquitoes, is dependent upon a complex interaction between the virus and vector [2, 6]. The virus in the ingested bloodmeal has to infect and replicate in the epithelial cells of the midgut (midgut infection barrier-MIB). The virus must then successfully escape from the midgut (midgut escape barrier-MEB) and infect the salivary glands (gland infection barrier-SGIB), followed by release of the virus into the salivary ducts for transmission orally to vertebrates. Salivary gland infection and escape barriers (salivary gland escape barrier-SGEB) determine if the virus can replicate and shed into the mosquito’s saliva for final transmission to the vertebrate host during *2, 7+. Mosquito vector capacity [susceptibility, extrinsic incubation period (EIP), transmission proportions], longevity, and host bloodmeal preferences are included among intrinsic factors. Each of these factors is affected by extrinsic factors such as larval density and nutrition, temperature, rainfall, avian host availability and avian host immunity [2, 7].

In Argentina, the Eared Dove (*Zenaida auriculata*) and Picui Ground-Dove (*Columbina picui*) are natural hosts of SLEV, while *Culex pipiens quinquefasciatus* is an efficient vector for the virus [3, 8]. For example, SLEV was detected and isolated in *Cx. p. quinquefasciatus* mosquitoes during an outbreak of encephalitis in humans in Córdoba City [9]. Infected *Cx. p. quinquefasciatus* have been detected during periods without reports of clinical disease symptoms in Santa Fe province between 1978-1983 [10] and Córdoba between 2001-2004 [11]. In addition viral isolations from field collected mosquitoes, and the population abundance throughout the Córdoba City [9], laboratory studies have confirmed horizontal transmission [8, 12], and the infrequent vertical transmission of SLEV in *Cx. p. quinquefasciatus* [13]. The presence of naturally SLEV-infected *Culex interfor* and *Culex saltanensis*, dominant species in urban-vegetated sub-assemblage that frequently feed on competent hosts, suggest they could participate as vectors in the transmission network of SLEV [3, 8, 9, 14]. Based on these evidences we argue that *Cx. saltanensis* and *Cx. interfor* could be playing a role as vectors of SLEV in central Argentina; yet vector competence experiments have not been conducted on these species. Therefore, we evaluated the vector competence of *Cx. interfor* and *Cx. saltanensis* against SLEV from central Argentina compared to the primary urban vector, *Cx. p. quinquefasciatus*.

## Methods

### Capture and maintenance of mosquito colonies

Egg rafts of *Cx. p. quinquefasciatus, Cx. interfor* and *Cx. saltanensis* were collected during 2015 at the Bajo Grande sewage treatment plant (31°24’13”S 64°06’08”W) located in east of Córdoba City. The site is surrounded by aquatic vegetation, reservoirs, low income human settlements and crop lands (vegetables and fruits). Authorization for mosquito field collections was obtained from the Municipality of Córdoba.

Egg rafts were collected with a dipper (350 ml) and transferred to polypropylene bottles with a fine brush and transported to the Instituto de Virología “Dr. *J. M. Vanella*” Facultad de Medicina, Universidad Nacional de Córdoba (FM-UNC). Mosquitoes were maintained 27°C, 70% humidity, and 12:12 h light: dark (L: D) photoperiodic. Adult F1 females from the field collected rafts were used in the vector competence assay (Table 1). Egg rafts were placed in a plastic container with distilled water and hatched larvae were fed with suspension of liver powder (0.25mg/0.5ml distilled water) every 48 h. After identification, larvae were grouped by species, pupae of the same species were placed inside screened cages (26 × 22 cm) until emergence, and adults provided a 10% sugar solution by soaked cotton balls. Identification of adult females and 4th instars were based on morphological keys [15]. Adult males were also collected to confirm the taxonomic identification based on the genitalia morphology [16, 17]. The small number of adult females used in this study, was caused by low feeding success and high mortality, which limited the sample size (Table 1).

**Table 1:**
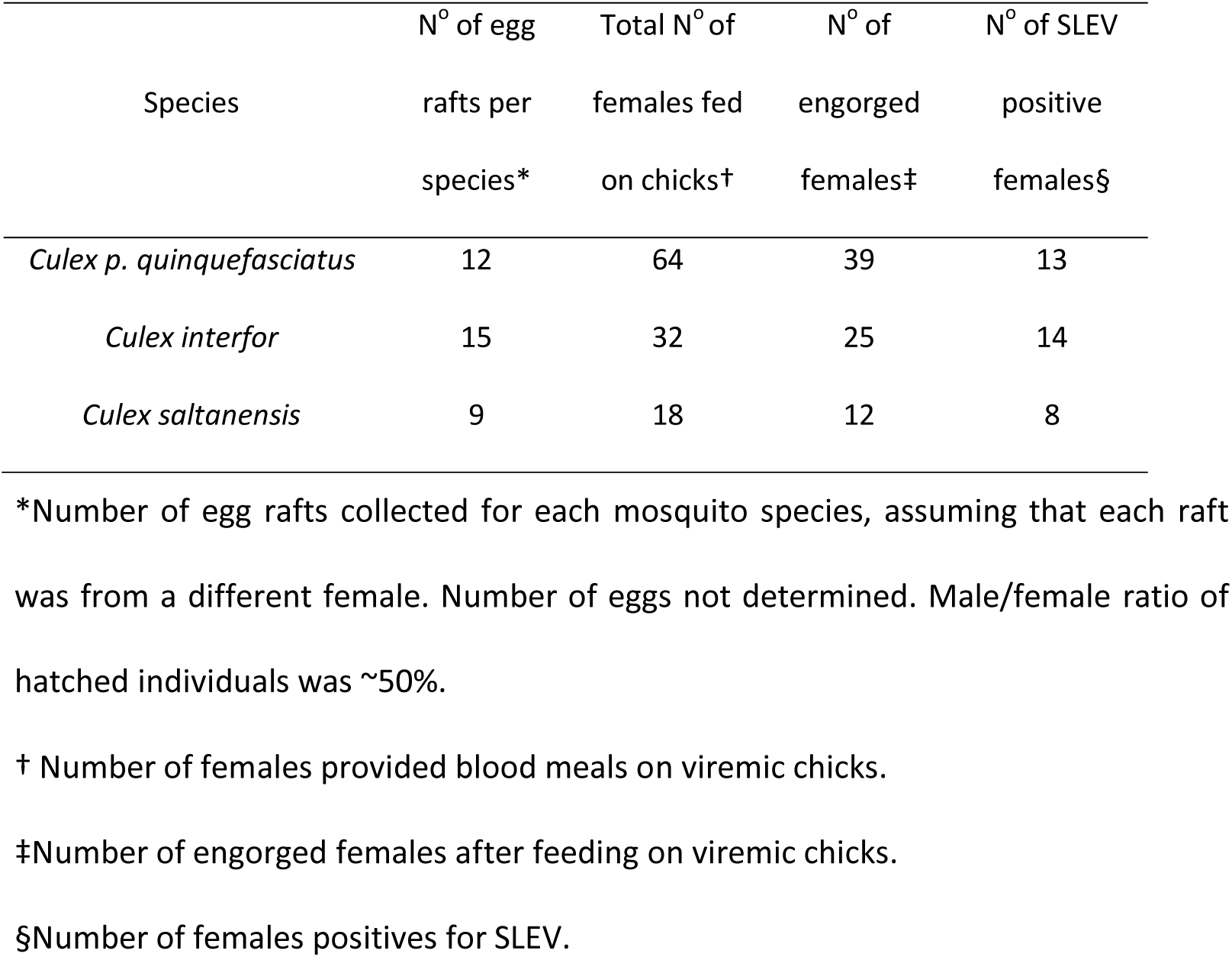
The egg collections and number females of Cx. p. quinquefasciatus, Cx. interfor and Cx. saltanensis in Córdoba City.

### Viral stock

Adult female mosquitoes were orally infected by feeding on viremic chicks that had been inoculated with SLEV CbaAr-4005 [9]. Viral stock was obtained from collections stored at the Instituto de Virología “Dr. J. M. Vanella” (FM-UNC). Viral stock was prepared from brain of an infected Swiss albino suckling mice homogenized in 10% P/V solution in Eagle’s minimal essential medium (MEM) (Gibco, Ireland), supplemented with 10% fetal bovine serum (FBS) (Natocor, Argentina) and 1% of gentamycin (Klonal, Argentina).

Viral titration was carried out by viral plaque assay in VERO cell monolayer (African green monkey kidney, Cercophitecus aethiops). We inoculated 0.1 ml of the samples onto VERO cell monolayer on a 12-well plate and incubated the plates for 60 min at 37°C with 5% CO2 and a humid atmosphere to favor the adsorption of the virus to the cell. After incubation 0.5 ml of MEM 2x was mixed with 1% methylcellulose and added to plate followed by incubation at 37°C for 7 days, until the formation of lytic plaques. The plates were then fixed with 10% formalin solution for 2 h and stained with crystal violet for the observation and plaque forming units (PFU) counts. Viral concentrations were then expressed as log10 PFU per milliliter (PFU/ml).

### Vector competence assay

Twenty-four hour-old chicks (Gallus gallus) were inoculated intraperitoneally with 0.1 ml of a viral suspension containing approximately 400 PFU of SLEV. In accordance with viremia kinetic [8], 48 h post-inoculation starved female mosquitoes were fed on chicks (Table 1). Prior to and following blood feeding by mosquitoes, 0.1 ml of blood was taken from the chick jugular vein to determine the viremia titer (potential viral load ingested by mosquitoes). The chick blood was diluted in 0.45 ml of MEM supplemented with 10% FBS and 1% gentamycin, followed by centrifugation at 4°C for 20 min at 2,300 g before supernatant stored at −80°C. Titers were determined by plaque assay and expressed as the mean viremia load before and after feeding (log10 PFU/ml). Mosquitoes were anesthetized by cold and fully engorged females were maintained in a screened cage at 27oC, H° 70%, 12:12 h L: D photoperiod and provided a 10% sugar solution. After 14 days EIP, the females were aspirated and anesthetized for 2 min at 4oC, placed on a refrigerated plate where the legs and wings were gently removed. Saliva samples were recovered after live females were placed on a flat surface with adhesive tape and the proboscis inserted for 30 min into a capillary tube with 0.001 ml glycerin (Todo Droga, Argentina) [18]. The abdomen was then removed, and saliva, legs and abdomen samples individually stored at-80oC with 1 ml of MEM supplemented with 10% FBS and 1% gentamycin. Legs and abdomen were individually homogenized by agitation with glass beads for 4 min and centrifuged at 11,200 g for 30 min. Subsequently, 0.1 ml of the sample was inoculated with two cellular and viral controls and infective virus particles detected by plaque assay as described above.

SLEV-negative saliva samples from females with positive abdomen samples were re-analyzed to reduce false negatives due to dilution. Thus, 0.2 ml of each sample was inoculated in two cell monolayer plates (‘A’ and ‘B’ plates) using the same controls as mentioned above. ‘A’ plates followed the same culture protocol described above. For ‘B’ plates, MEM, FBS (2%) and gentamycin (1%) were added, and after 96 h the cytopathic effect (cell detachment, rounding and nonconfluent monolayer) was determined under an inverted microscope. To amplify viral particles not detected in the first passage, monolayer from the ‘B’ plates were harvested on the fourth day post inoculation, by collecting 0.75 ml from each of the duplicates wells followed by centrifugation at 9,300 g for 30 min (passage 1). Finally, another plate was inoculated (duplicate, same ‘B’ plates protocol) with 0.2 ml of the samples without cytopathic effect in the ‘B’ plates. Therefore, PFU/ml was recorded in ‘A’ plates and the cytopathic effect presence in ‘B’ plates.

### Ethical Guidelines

The protocol used was approved by Consejo Nacional de Investigaciones Científicas y Técnicas (CONICET) and Instituto de Virología “Dr. J. M. Vanella” (FM-UNC). The chicks were maintained in cages with a substrate of wood shavings and with a 12:12 h L: D photoperiod, 40-80% humidity and 20-26°C temperature. Food and water were available ad libitum (balanced concentrate feed, Gepsa Feeds, Argentina). The experimental procedures used were to minimize or eliminate pain and distress. Awareness were taken to avoid chicks suffering, as anesthetic protocols were not used. Euthanasia was carried out with neck dislocation by a trained laboratory technician.

### Statistics analysis

Abdomens, legs, and saliva were considered positive when they showed at least one PFU or the presence of cytopathic effect on the cell culture monolayer. Infection rates were defined as the numbers of positive mosquito abdomens of the total number of blood feed mosquitoes. Dissemination rates were calculated as the number of mosquitoes with positive legs out of the number of mosquitoes fed analyzed. Transmission rates were determined by the number of mosquitoes with positive saliva of the total number of mosquitoes that blood fed. Confidence intervals (0.95%) for infection, dissemination and transmission rates and graphical presentations were made in the R [19]. Differences in rates for each species Culex spp. were compared by a Fisher exact test, considering statistically significant α < 0.05 [20].

## Results

The numbers of eggs collected *Cx. p. quinquefasciatus, Cx. interfor, Cx. saltanensis* and females that fed and were positive for SLEV are shown in Table 1. Mosquitoes were permissive to orally acquire infections, to virus dissemination, and transmission of SLEV in saliva (Figure 1). There was a narrow range of viremia during blood feeding (*Cx. p. quinquefasciatus* = 2.9 log_10_ PFU/ml, *Cx. interfor* = 3.5 log_10_ PFU/ml and *Cx. saltanensis* = 3.2 log_10_ PFU/ml).

**Figure 1:**
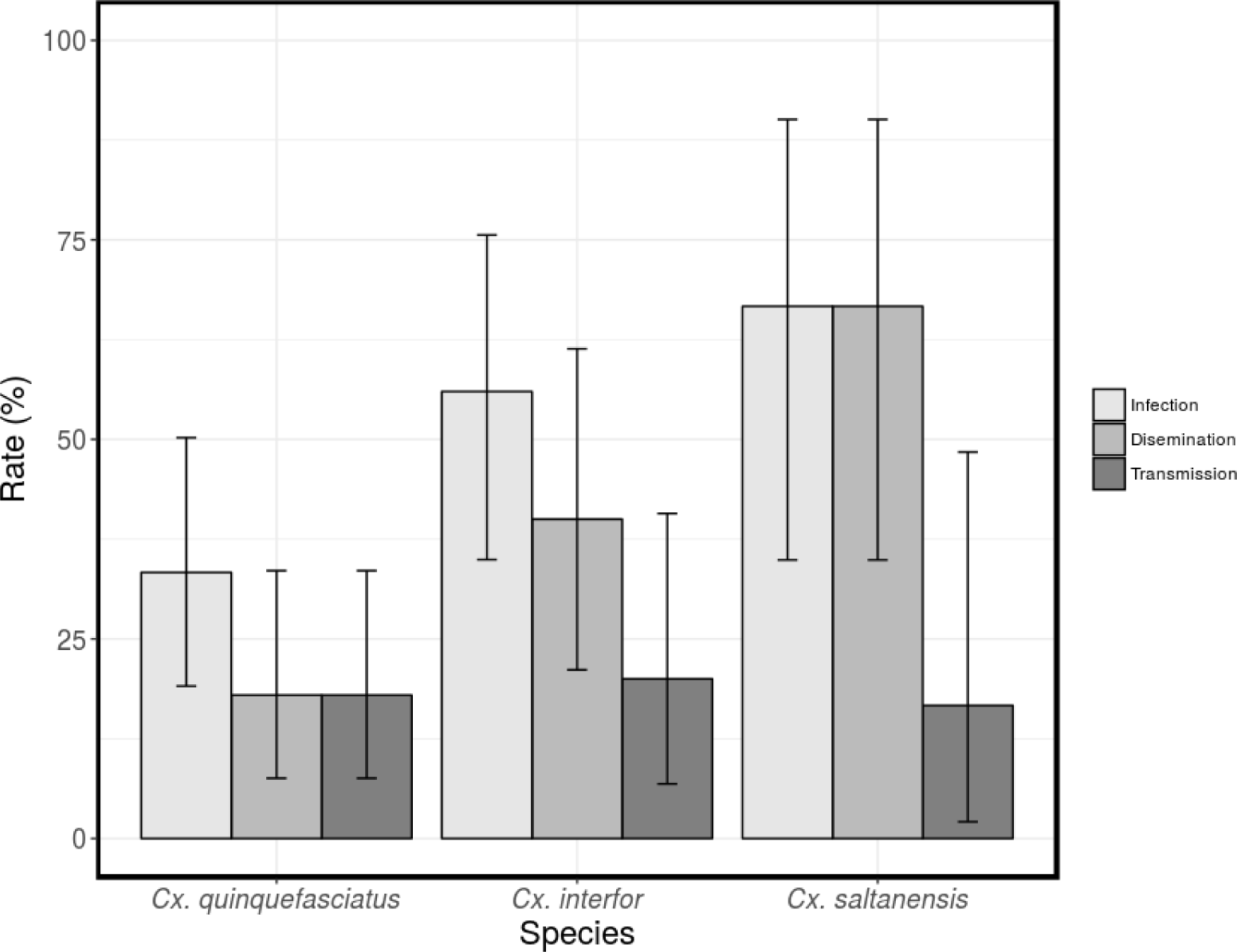
Vector competence for SLEV of *Cx. p. quinquefasciatus, Cx. interfor* and *Cx. saltanensis*. Infection rate (number of positive mosquito abdomens/total number of mosquitoes fed) in light gray, dissemination rate (number of mosquitoes with positive legs/total number of mosquitoes fed) in gray and transmission rate (number of mosquitoes with positive saliva/total number of mosquitoes fed) in dark gray; with their 0.95% CIs.

Females *Cx. p. quinquefasciatus* were equally susceptible to infection and transmission of SLEV, because were not statistically significant difference between infection (13/39, 33%), dissemination (7/13, 54%) and transmission rates (18%, 7/39) (Table 2).

**Table 2:** Vector competence of *Culex p. quinquefasciatus, Cx. interfor* and *Cx. saltanensis* for SLEV measured as infection, dissemination and transmission.

| Species<br>Vector<br>competence | *Culex p. quinquefasciatus* |  |  | *Culex interfor* |  |  | *Culex saltanensis* |  |  |
| --- | --- | --- | --- | --- | --- | --- | --- | --- | --- |
|  | N* | Rates†<br>(0.95%CI) | Viral<br>load‡<br>(0.95%CI) | N* | Rates†<br>(0.95%CI) | Viral<br>load‡<br>(0.95%CI) | N* | Rates†<br>(0.95%CI) | Viral load‡<br>(0.95%CI) |
| Infection | 13/39 | 33<br>(19.1-50.2) | 2.4<br>(1.3-2.9) | 14/25 | 56<br>(34.9-75.6) | 4.8<br>(2.4-5.3) | 8/12 | 67<br>(34.9-90.1) | 4.8<br>(3.8-5.1) |
| Dissemination | 7/39 | 18<br>(7.5-33.5) | 2.9 | 10/25 | 40<br>(21.1-61.3) | 4.8<br>(1-5.4) | 8/12 | 67<br>(34.9-90.1) | 4.6<br>(1.6-5.1) |
| Transmission | 7/39 | 18<br>(7.5-33.5) | No data | 5/25 | 20<br>(6.8-40.7) | 1.8<br>(1.1-2.3) | 2/12 | 17<br>(2.1-48.4) | 1.8 |

\*Number of mosquitoes positive for SLEV/total number of mosquitoes assayed
†Infection rate (number of positive mosquito abdomens/total number of mosquitoes fed); dissemination rate (number of mosquitoes with positive legs/total number of mosquitoes fed) and transmission rate (number of mosquitoes with positive saliva/total number of mosquitoes fed).
‡ Average SLEV titers ( $\log_{10}$ PFU/ml) in abdomen, legs and saliva of each mosquito species.

Viral loads were evaluated for abdomens 2.4 log_10_ PFU/ml and legs 2.9 log_10_ PFU/ml. Because lysis plaques in *Cx. p. quinquefasciatus* were not confluent the SLEV load in saliva was not able to measure (Table 2). For *Cx. interfor*, statistically significant differences were observed across infection (14/25, 56%) and transmission rates (20%, 5/25) (Fisher’s exact test, p = 0.0186) (Table 3).

**Table 3:**
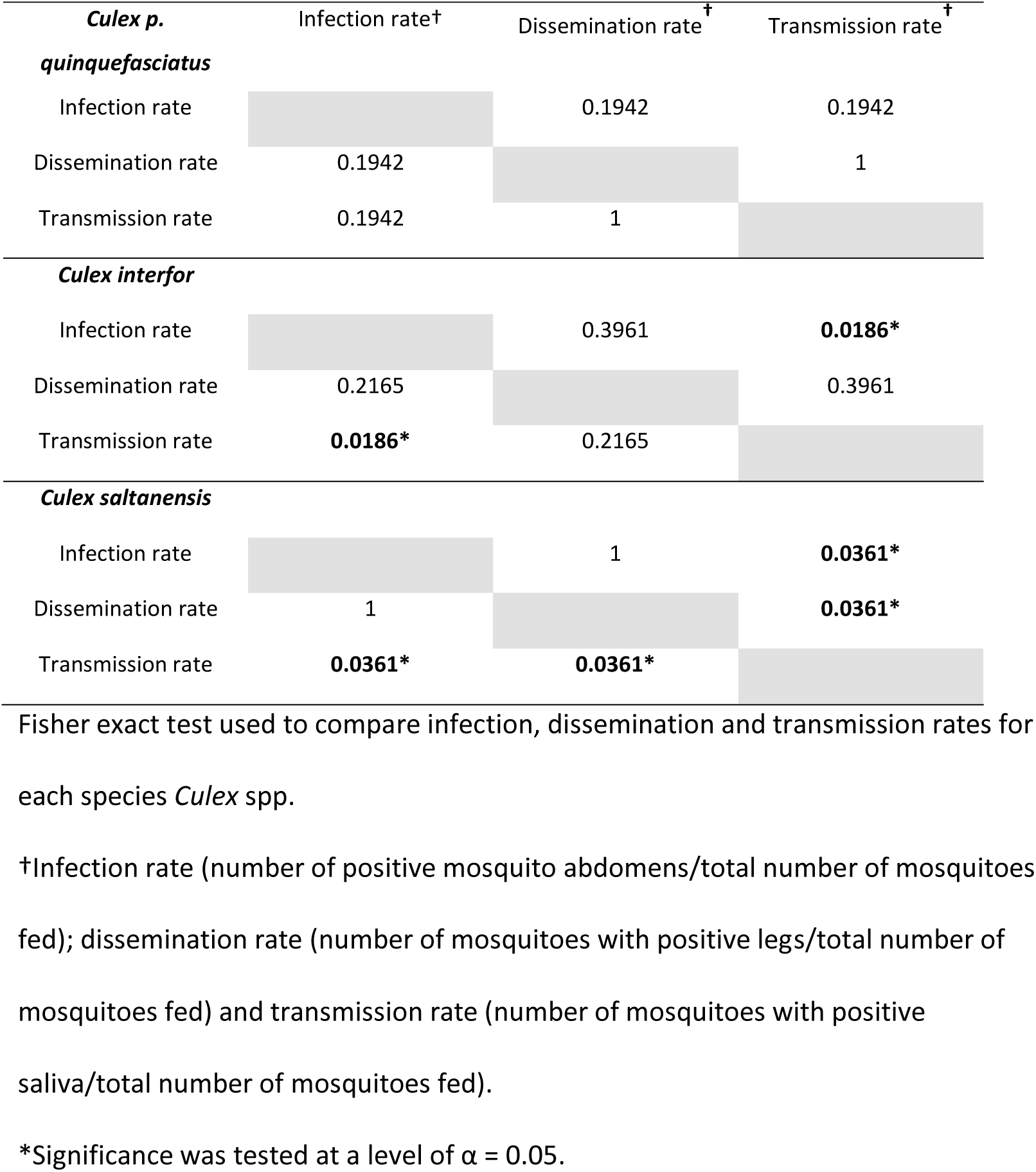
Infection, dissemination and transmission rates of *Cx. p. quinquefasciatus, Cx. interfor* and *Cx. saltanensis* in Córdoba City.

Viral loads were evaluated for abdomens 4.8 log_10_ PFU/ml, legs 4.8 log_10_ PFU/ml and saliva 1.8 log_10_ PFU/ml (Table 2). For *Cx. saltanensis*, statistically significant differences were observed between infection (8/12, 67%) and transmission rates (17%, 2/12), and this rate with respect to the dissemination rate (8/12, 67%) (Fisher’s exact test, p = 0.0361) (Table 3). Viral loads were evaluated for abdomens 4.8 log_10_ PFU/ml, legs 4.6 log_10_ PFU/ml and saliva 1.8 log_10_ PFU/ml (Table 2).

## Discussion

*Culex p. quinquefasciatus, Cx. interfor* and *Cx. saltanensis* were competent vectors for SLEV based on 1) acquired infections, 2) disseminated virus, and 3) transmission of SLEV in the saliva after feeding on a viremic chick. SLEV was able to cross the midgut barriers showing disseminated infection (positive legs) and was able to cross salivary gland barriers in positive saliva samples. However, the number of tested mosquitoes was low, thus our results are not conclusive.

To be considered a vector, a mosquito species must fulfill several biological and ecological characteristics [21, 22]. The coevolution between pathogen and arthropods determine the vector competence, and thus the ability to acquire, maintain and eventually transmit it [21]. Variation in vector competence has been documented with all of the major disease agents they transmit (i.e. malaria and filarial parasites, and arboviruses) [6, 23, 24]. Although the life cycle of each pathogen is distinct, they all face the common events of being ingested, exposed to the midgut environment, and traversing hemocoel to reach their tissue site of development and/or suitable site for transmission back to a new vertebrate host. Each of these migratory steps presents potential barriers that might be manipulated to interfere with normal pathogen migration and/or development [22]. Along with these barriers, some other factors like digestive enzymes, midgut microbiota, and innate immune responses might be responsible for vector’s refractoriness and ineffective horizontal transmission [2]. Understanding the vector competence is crucial for assessing the risks of arbovirus transmission and maintenance in nature.

There is considerable specificity in the vector-arbovirus relationship, and some of this specificity comes from the ability of a particular arbovirus to overcome tissue barriers in the vector to establish a persistent infection. Factors that strongly affect vector competence of a mosquito for a particular arbovirus include MIB, MEB, SGIB, and SGEB *6+. In our findings, among infected mosquitoes, dissemination was achieved in 100% of those individuals tested (8/8) for *Cx. saltanensis*, while are 71% (10/14) of *Cx. interfor* and 54% (7/13) of *Cx. p. quinquefasciatus* demonstrated disseminated infections. These results suggest the possible existence of a midgut barrier to SLEV in *Cx. p. quinquefasciatus*. Kramer et al. were the first to demonstrate that the inability of infected *Cx. tarsalis* mosquitoes to transmit western equine encephalitis virus (WEEV) was associated with a MEB [25]. Also following studies detected existence the midgut barrier for Rift Valley Fever virus in *Cx. pipiens* [26], SLEV and WNV in *Cx. p. quinquefasciatus* respectively [27, 28].

Virus transmission is a critical component of laboratory studies of vector competence and is essential to understanding the epidemiology of arboviruses [18]. Reisen et al. quantified the viral particles of SLEV expectorated in the saliva of *Cx. tarsalis* (1.1-2.2 log_10_ PFU) [29] and we obtained similar data for *Cx. interfor* (range = 1.1-2.3 log_10_ PFU) and *Cx. saltanensis* (1.8 log_10_ PFU). Moreover, all *Cx. p. quinquefasciatus* females with disseminated infections demonstrated SLEV in their saliva (7/7); this rate was only 50% (5/10) in *Cx. interfor* and 25% (2/8) in *Cx. saltanensis*, indicating a potential salivary gland barrier in both *Cx. interfor* and *Cx. saltanensis*. Similar results were obtained for Japanese encephalitis virus in *Cx. p. molesus* [30], WEEV in *Cx. tarsalis* [25] and Venezuelan equine encephalitis virus in *Psorophora cingulata* and *Coquillettidia venezuelensis* [31]. Further work are needed should evaluate and explore the relationship between midgut and salivary gland barriers.

Our results corroborate the findings reported by Diaz et al. on the susceptibility of *Cx. p. quinquefasciatus* for SLEV infection [8]. These authors observed an infection rate of 70% using the same viral strain CbaAr-4005 and feeding viremia level of 5.2 log_10_ PFU/ml. In our study only one third of the *Cx. p. quinquefasciatus* females became infected. This difference could be related to the lower viremia level that the mosquitoes were exposed to in this assay (2.9 log_10_ PFU/ml). Mitchell et al. obtained a transmission rate of (90.5%, 19/21) with strain 78V-6507 for *Cx. p. quinquefasciatus* from Santa Fe Province [12]. In our study, the transmission rate was (18%, 7/39). Dissemination and transmission rate formulation in Mitchell and herein are different. Even through dissemination rates are higher with a higher infectious dose (Mitchell, 4.1-4.8 log_10_ PFU/ml vs our study, 2.9 log_10_ PFU/ml). The effect of viremia could be affected by the small number of individuals used in this study. Viremia, virus dose, extrinsic incubation temperature, mosquito age, and colony are all important factors influencing the vector competence of *Cx. p. quinquefasciatus* [27, 32]. Future studies to determine the Minimum Infection Threshold and the Extrinsic Incubation Periods are needed.

The maintenance of SLEV in nature is complex and requires the coexistence in time and space of mosquito vectors and avian hosts. The detection of naturally infected mosquitoes does not represent, by itself, a reliable proof of their role as a competent vector. In the case of *Cx. p. quinquefasciatus, Cx. interfor* and *Cx. saltanensis*, there is evidence supporting their intervention as vectors in SLEV transmission. *Culex p. quinquefasciatus* is considered the primary vector, because it was abundant and *pools* assayed were positive for SLEV [9-11, 14]. *Culex saltanensis* and *Cx. interfor* participate in the maintenance of SLEV and could assist in the spillover of SLEV to humans. In 2004, prior to the outbreak of encephalitis in Córdoba City, SLEV infected *Cx. interfor* were detected [11]. In 2010, there were small outbreaks of SLEV in provinces, e.g., Buenos Aires, Córdoba and San Juan [33]. Not long thereafter, SLEV infected *Cx. saltanensis* were detected for first time in Córdoba City [14]. *Culex interfor* and *Cx. saltanensis* are mainly ornithophiles, and bloodmeal from Columbiformes and Passeriformes have been also detected, although the pattern of host preference and its drivers have not been established yet [14, 34, 35]. In Argentina, *Z. auriculata* and *C. picui* are amplifier hosts of SLEV and have been recorded in engorged *Cx. saltanensis, Cx. interfor* and *Cx. p. quinquefasciatus* sustaining that SLEV maintenance could relied on multiple vectors [3, 36]. However, the transmission load in SLEV episystem could be unequal between the three *Culex*, despite they showed similar transmission rate experimentally (ranged 17-20%). For instance, among other traits, lifespan difference among mosquito species is expected to impact vector capacity as longest lifespan increase the odds of extrinsic incubation completeness and delivering infectious bites [7, 21]. Here, *Cx. p. quinquefasciatus* was less able to survive after a viremic bloodmeal than *Cx. saltanensis* and *Cx. interfor* suggesting that the role of the last species has been neglected. In addition, it has been proposed that *Cx. interfor* could transmit SLEV from birds to mammals and thus fulfill a role of “bridge vector” *14, 34, 37+ as *Cx. interfor* was recorded in human baited barley traps [37] and *Cx. interfor* and *Cx. saltanensis* switch between bird feeding profile in spring-summer to bird-mammals in autumn in a rural environment [38]. The local populations of *Culex* spp. increase in abundance with peaks in summer, with are temporal distribution of *Culex* spp. coinciding with the activity peaks of SLEV in human infection [3, 39]. Adult mosquitoes belonging to the species *Cx. saltanensis* and *Cx. interfor* have been found in increasing numbers in Córdoba City, with a higher abundance in urban and periurban areas where vegetation is more robust. This differs from *Cx. p. quinquefasciatus*, which is predominant throughout a vast range of city-type environments [39]. Our results support the hypothesis that SLEV is transmitted by multiple sympatric *Culex* spp., and that both *Cx. saltanensis* and *Cx. interfor* can be considered potential vectors of SLEV. In the United States, this has been observed as well; however, different mosquito species serve as the primary vector transmitting SLEV in different geographical areas. *Culex quinquefasciatus* and *Cx. nigripalpus* are vectors for the virus in Florida, *Cx. tarsalis* in the western and *Cx. pipiens* in the northern United States [4].

SLEV is a multi-host and multi-vector flavivirus in the process of an ongoing reemergence in Argentina. Further studies are required to understand the spatial compartmentalization of these mosquito species in the transmission network of SLEV by performing vector capacity studies. By having insights into its ecoepidemiology, we will have a better understanding of which factors are causing this reemergence and how biological and environmental factors interact with and affect its activity. In addition, knowledge of the potential mosquito species vectors of SLEV will provide information to be used by different public agencies related to human health for the control of vector mosquito populations and improve efficiency in SLEV prevention programs.

## Data accessibility

Data are available online: link or DOI of the webpage hosting the data.

## Supplementary material

End of the Manuscript

## Acknowledgements

We thank Kristin Sloyer for sharing suggestions and improving the English of the manuscript.

Version 6 of this preprint has been peer-reviewed and recommended by Peer Community In Entomology (https://doi.org/10.24072/pci.entomol.100002).

## Conflict of interest disclosure

The authors of this preprint declare that they have no financial conflict of interest with the content of this article.

Adrian Diaz is one of the PCI Entomology recommenders.

## Supplementary material

**Table:** Female mosquitoes *Cx. interfor, Cx. saltanensis* and *Cx. p. quinquefasciatus* used in vector competence assay with the SLEV

| Specie | ID | Viremia | Results | Rate | N | PFU/ml | log<sub>10</sub>PFU/ml |
| --- | --- | --- | --- | --- | --- | --- | --- |
| *Cx.interfor* | 1.1 | 3.5 | 0 | Infection | 25 | 0 |  |
| *Cx.interfor* | 1.1 | 3.5 | 0 | Dissemination | 25 | 0 |  |
| *Cx.interfor* | 1.1 | 3.5 | 0 | Transmission | 25 | 0 |  |
| *Cx.interfor* | 1.2 | 3.5 | 0 | Infection | 25 | 0 |  |
| *Cx.interfor* | 1.2 | 3.5 | 0 | Dissemination | 25 | 0 |  |
| *Cx.interfor* | 1.2 | 3.5 | 0 | Transmission | 25 | 0 |  |
| *Cx.interfor* | 1.3 | 3.5 | 0 | Infection | 25 | 0 |  |
| *Cx.interfor* | 1.3 | 3.5 | 0 | Dissemination | 25 | 0 |  |
| *Cx.interfor* | 1.3 | 3.5 | 0 | Transmission | 25 | 0 |  |
| *Cx.interfor* | 1.4 | 3.5 | 0 | Infection | 25 | 0 |  |
| *Cx.interfor* | 1.4 | 3.5 | 0 | Dissemination | 25 | 0 |  |
| *Cx.interfor* | 1.4 | 3.5 | 0 | Transmission | 25 | 0 |  |
| *Cx.interfor* | 1.5 | 3.5 | 0 | Infection | 25 | 0 |  |
| *Cx.interfor* | 1.5 | 3.5 | 0 | Dissemination | 25 | 0 |  |
| *Cx.interfor* | 1.5 | 3.5 | 0 | Transmission | 25 | 0 |  |
| *Cx.interfor* | 1.6 | 3.5 | 1 | Infection | 25 | 145000 | 5,161368002 |
| *Cx.interfor* | 1.6 | 3.5 | 1 | Dissemination | 25 | 82000 | 4,913813852 |
| *Cx.interfor* | 1.6 | 3.5 | 1 | Transmission | 25 | 12,5 | 1,096910013 |
| *Cx.interfor* | 1.7 | 3.5 | 1 | Infection | 25 | 750 | 2,875061263 |
| *Cx.interfor* | 1.7 | 3.5 | 0 | Dissemination | 25 | 0 |  |
| *Cx.interfor* | 1.7 | 3.5 | 0 | Transmission | 25 | 0 |  |
| *Cx.interfor* | 1.8 | 3.5 | 1 | Infection | 25 | 1320 | 3,120573931 |
| *Cx.interfor* | 1.8 | 3.5 | 0 | Dissemination | 25 | 0 |  |
| *Cx.interfor* | 1.8 | 3.5 | 0 | Transmission | 25 | 0 |  |
| *Cx.interfor* | 1.9 | 3.5 | 1 | Infection | 25 | 280 | 2,447158031 |
| *Cx.interfor* | 1.9 | 3.5 | 0 | Dissemination | 25 | 0 |  |
| *Cx.interfor* | 1.9 | 3.5 | 0 | Transmission | 25 | 0 |  |
| *Cx.interfor* | 1.10 | 3.5 | 0 | Infection | 25 | 0 |  |
| *Cx.interfor* | 1.10 | 3.5 | 0 | Dissemination | 25 | 0 |  |
| *Cx.interfor* | 1.10 | 3.5 | 0 | Transmission | 25 | 0 |  |
| *Cx.interfor* | 1.11 | 3.5 | 0 | Infection | 25 | 0 |  |
| *Cx.interfor* | 1.11 | 3.5 | 0 | Dissemination | 25 | 0 |  |
| *Cx.interfor* | 1.11 | 3.5 | 0 | Transmission | 25 | 0 |  |
| *Cx.interfor* | 1.12 | 3.5 | 1 | Infection | 25 | 140000 | 5,146128036 |
| *Cx.interfor* | 1.12 | 3.5 | 1 | Dissemination | 25 | 230000 | 5,361727836 |
| *Cx.interfor* | 1.12 | 3.5 | 0 | Transmission | 25 | 0 |  |
| *Cx.interfor* | 1.13 | 3.5 | 0 | Infection | 25 | 0 |  |
| *Cx.interfor* | 1.13 | 3.5 | 0 | Dissemination | 25 | 0 |  |
| *Cx.interfor* | 1.13 | 3.5 | 0 | Transmission | 25 | 0 |  |
| *Cx.interfor* | 1.14 | 3.5 | 0 | Infection | 25 | 0 |  |
| *Cx.interfor* | 1.14 | 3.5 | 0 | Dissemination | 25 | 0 |  |
| *Cx.interfor* | 1.14 | 3.5 | 0 | Transmission | 25 | 0 |  |
| *Cx.interfor* | 1.15 | 3.5 | 1 | Infection | 25 | 182000 | 5,260071388 |
| *Cx.interfor* | 1.15 | 3.5 | 1 | Dissemination | 25 | 23000 | 4,361727836 |
| *Cx.interfor* | 1.15 | 3.5 | 1 | Transmission | 25 | 212,5 | 2,327358934 |
| *Cx.interfor* | 1.16 | 3.5 | 0 | Infection | 25 | 0 |  |
| *Cx.interfor* | 1.16 | 3.5 | 0 | Dissemination | 25 | 0 |  |
| *Cx.interfor* | 1.16 | 3.5 | 0 | Transmission | 25 | 0 |  |
| *Cx.interfor* | 1.17 | 3.5 | 1 | Infection | 25 | 116000 | 5,064457989 |
| *Cx.interfor* | 1.17 | 3.5 | 1 | Dissemination | 25 | 20000 | 4,301029996 |
| *Cx.interfor* | 1.17 | 3.5 | 1 | Transmission | 25 | 20 | 1,301029996 |
| *Cx.interfor* | 1.18 | 3.5 | 1 | Infection | 25 | 127000 | 5,103803721 |
| *Cx.interfor* | 1.18 | 3.5 | 1 | Dissemination | 25 | 13000 | 4,113943352 |
| *Cx.interfor* | 1.18 | 3.5 | 1 | Transmission | 25 | 20 | 1,301029996 |
| *Cx.interfor* | 1.19 | 3.5 | 1 | Infection | 25 | 960 | 2,982271233 |
| *Cx.interfor* | 1.19 | 3.5 | 1 | Dissemination | 25 | 10 | 1 |
| *Cx.interfor* | 1.19 | 3.5 | 0 | Transmission | 25 | 0 |  |
| *Cx.interfor* | 1.20 | 3.5 | 1 | Infection | 25 | 75000 | 4,875061263 |
| *Cx.interfor* | 1.20 | 3.5 | 1 | Dissemination | 25 | 60000 | 4,77815125 |
| *Cx.interfor* | 1.20 | 3.5 | 0 | Transmission | 25 | 0 |  |
| *Cx.interfor* | 1.21 | 3.5 | 1 | Infection | 25 | 930 | 2,968482949 |
| *Cx.interfor* | 1.21 | 3.5 | 1 | Dissemination | 25 | 150 | 2,176091259 |
| *Cx.interfor* | 1.21 | 3.5 | 1 | Transmission | 25 | NA |  |
| *Cx.interfor* | 1.22 | 3.5 | 1 | Infection | 25 | 20000 | 4,301029996 |
| *Cx.interfor* | 1.22 | 3.5 | 0 | Dissemination | 25 | 0 |  |
| *Cx.interfor* | 1.22 | 3.5 | 0 | Transmission | 25 | 0 |  |
| *Cx.interfor* | 1.23 | 3.5 | 0 | Infection | 25 | 0 |  |
| *Cx.interfor* | 1.23 | 3.5 | 0 | Dissemination | 25 | 0 |  |
| *Cx.interfor* | 1.23 | 3.5 | 0 | Transmission | 25 | 0 |  |
| *Cx.interfor* | 1.24 | 3.5 | 1 | Infection | 25 | 180000 | 5,255272505 |
| *Cx.interfor* | 1.24 | 3.5 | 1 | Dissemination | 25 | 80000 | 4,903089987 |
| *Cx.interfor* | 1.24 | 3.5 | 0 | Transmission | 25 | 0 |  |
| *Cx.interfor* | 1.25 | 3.5 | 1 | Infection | 25 | 1050 | 3,021189299 |
| *Cx.interfor* | 1.25 | 3.5 | 1 | Dissemination | 25 | 69000 | 4,838849091 |
| *Cx.interfor* | 1.25 | 3.5 | 0 | Transmission | 25 | 0 |  |
| *Cx.saltanensis* | 1.1 | 3.2 | 1 | Infection | 12 | 43000 | 4,633468456 |
| *Cx.saltanensis* | 1.1 | 3.2 | 1 | Dissemination | 12 | 40 | 1,602059991 |
| *Cx.saltanensis* | 1.1 | 3.2 | 1 | Transmission | 12 | NA |  |
| *Cx.saltanensis* | 1.2 | 3.2 | 1 | Infection | 12 | 88000 | 4,944482672 |
| *Cx.saltanensis* | 1.2 | 3.2 | 1 | Dissemination | 12 | 70000 | 4,84509804 |
| *Cx.saltanensis* | 1.2 | 3.2 | 0 | Transmission | 12 | 0 |  |
| *Cx.saltanensis* | 1.3 | 3.2 | 1 | Infection | 12 | 74000 | 4,86923172 |
| *Cx.saltanensis* | 1.3 | 3.2 | 1 | Dissemination | 12 | 132000 | 5,120573931 |
| *Cx.saltanensis* | 1.3 | 3.2 | 1 | Transmission | 12 | 65 | 1,812913357 |
| *Cx.saltanensis* | 1.4 | 3.2 | 0 | Infection | 12 | 0 |  |
| *Cx.saltanensis* | 1.4 | 3.2 | 0 | Dissemination | 12 | 0 |  |
| *Cx.saltanensis* | 1.4 | 3.2 | 0 | Transmission | 12 | 0 |  |
| *Cx.saltanensis* | 1.5 | 3.2 | 0 | Infection | 12 | 0 |  |
| *Cx.saltanensis* | 1.5 | 3.2 | 0 | Dissemination | 12 | 0 |  |
| *Cx.saltanensis* | 1.5 | 3.2 | 0 | Transmission | 12 | 0 |  |
| *Cx.saltanensis* | 1.6 | 3.2 | 1 | Infection | 12 | 120000 | 5,079181246 |
| *Cx.saltanensis* | 1.6 | 3.2 | 1 | Dissemination | 12 | 45000 | 4,653212514 |
| *Cx.saltanensis* | 1.6 | 3.2 | 0 | Transmission | 12 | 0 |  |
| *Cx.saltanensis* | 1.7 | 3.2 | 0 | Infection | 12 | 0 |  |
| *Cx.saltanensis* | 1.7 | 3.2 | 0 | Dissemination | 12 | 0 |  |
| *Cx.saltanensis* | 1.7 | 3.2 | 0 | Transmission | 12 | 0 |  |
| *Cx.saltanensis* | 1.8 | 3.2 | 1 | Infection | 12 | 73000 | 4,86332286 |
| *Cx.saltanensis* | 1.8 | 3.2 | 1 | Dissemination | 12 | 1040 | 3,017033339 |
| *Cx.saltanensis* | 1.8 | 3.2 | 0 | Transmission | 12 | 0 |  |
| *Cx.saltanensis* | 1.9 | 3.2 | 1 | Infection | 12 | 7000 | 3,84509804 |
| *Cx.saltanensis* | 1.9 | 3.2 | 1 | Dissemination | 12 | 1650 | 3,217483944 |
| *Cx.saltanensis* | 1.9 | 3.2 | 0 | Transmission | 12 | 0 |  |
| *Cx.saltanensis* | 1.10 | 3.2 | 1 | Infection | 12 | 79000 | 4,897627091 |
| *Cx.saltanensis* | 1.10 | 3.2 | 1 | Dissemination | 12 | 1260 | 3,100370545 |
| *Cx.saltanensis* | 1.10 | 3.2 | 0 | Transmission | 12 | 0 |  |
| *Cx.saltanensis* | 1.11 | 3.2 | 0 | Infection | 12 | 0 |  |
| *Cx.saltanensis* | 1.11 | 3.2 | 0 | Dissemination | 12 | 0 |  |
| *Cx.saltanensis* | 1.11 | 3.2 | 0 | Transmission | 12 | 0 |  |
| *Cx.saltanensis* | 1.12 | 3.2 | 1 | Infection | 12 | 76000 | 4,880813592 |
| *Cx.saltanensis* | 1.12 | 3.2 | 1 | Dissemination | 12 | 37000 | 4,568201724 |
| *Cx.saltanensis* | 1.12 | 3.2 | 0 | Transmission | 12 | 0 |  |
| *Cx.p.quinquefasciatus* | 1.1 | 2.9 | 0 | Infection | 39 | 0 |  |
| *Cx.p.quinquefasciatus* | 1.1 | 2.9 | 0 | Dissemination | 39 | 0 |  |
| *Cx.p.quinquefasciatus* | 1.1 | 2.9 | 0 | Transmission | 39 | 0 |  |
| *Cx.p.quinquefasciatus* | 1.2 | 2.9 | 0 | Infection | 39 | 0 |  |
| *Cx.p.quinquefasciatus* | 1.2 | 2.9 | 0 | Dissemination | 39 | 0 |  |
| *Cx.p.quinquefasciatus* | 1.2 | 2.9 | 0 | Transmission | 39 | 0 |  |
| *Cx.p.quinquefasciatus* | 1.3 | 2.9 | 0 | Infection | 39 | 0 |  |
| *Cx.p.quinquefasciatus* | 1.3 | 2.9 | 0 | Dissemination | 39 | 0 |  |
| *Cx.p.quinquefasciatus* | 1.3 | 2.9 | 0 | Transmission | 39 | 0 |  |
| *Cx.p.quinquefasciatus* | 1.4 | 2.9 | 0 | Infection | 39 | 0 |  |
| *Cx.p.quinquefasciatus* | 1.4 | 2.9 | 0 | Dissemination | 39 | 0 |  |
| *Cx.p.quinquefasciatus* | 1.4 | 2.9 | 0 | Transmission | 39 | 0 |  |
| *Cx.p.quinquefasciatus* | 1.5 | 2.9 | 1 | Infection | 39 | 490 | 2,69019608 |
| *Cx.p.quinquefasciatus* | 1.5 | 2.9 | 0 | Dissemination | 39 | 0 |  |
| *Cx.p.quinquefasciatus* | 1.5 | 2.9 | 0 | Transmission | 39 | 0 |  |
| *Cx.p.quinquefasciatus* | 1.6 | 2.9 | 0 | Infection | 39 | 0 |  |
| *Cx.p.quinquefasciatus* | 1.6 | 2.9 | 0 | Dissemination | 39 | 0 |  |
| *Cx.p.quinquefasciatus* | 1.6 | 2.9 | 0 | Transmission | 39 | 0 |  |
| *Cx.p.quinquefasciatus* | 1.7 | 2.9 | 0 | Infection | 39 | 0 |  |
| *Cx.p.quinquefasciatus* | 1.7 | 2.9 | 0 | Dissemination | 39 | 0 |  |
| *Cx.p.quinquefasciatus* | 1.7 | 2.9 | 0 | Transmission | 39 | 0 |  |
| *Cx.p.quinquefasciatus* | 1.8 | 2.9 | 0 | Infection | 39 | 0 |  |
| *Cx.p.quinquefasciatus* | 1.8 | 2.9 | 0 | Dissemination | 39 | 0 |  |
| *Cx.p.quinquefasciatus* | 1.8 | 2.9 | 0 | Transmission | 39 | 0 |  |
| *Cx.p.quinquefasciatus* | 1.9 | 2.9 | 1 | Infection | 39 | NA |  |
| *Cx.p.quinquefasciatus* | 1.9 | 2.9 | 1 | Dissemination | 39 | NA |  |
| *Cx.p.quinquefasciatus* | 1.9 | 2.9 | 1 | Transmission | 39 | NA |  |
| *Cx.p.quinquefasciatus* | 1.10 | 2.9 | 0 | Infection | 39 | 0 |  |
| *Cx.p.quinquefasciatus* | 1.10 | 2.9 | 0 | Dissemination | 39 | 0 |  |
| *Cx.p.quinquefasciatus* | 1.10 | 2.9 | 0 | Transmission | 39 | 0 |  |
| *Cx.p.quinquefasciatus* | 1.11 | 2.9 | 1 | Infection | 39 | 370 | 2,568201724 |
| *Cx.p.quinquefasciatus* | 1.11 | 2.9 | 1 | Dissemination | 39 | NA |  |
| *Cx.p.quinquefasciatus* | 1.11 | 2.9 | 1 | Transmission | 39 | NA |  |
| *Cx.p.quinquefasciatus* | 1.12 | 2.9 | 0 | Infection | 39 | 0 |  |
| *Cx.p.quinquefasciatus* | 1.12 | 2.9 | 0 | Dissemination | 39 | 0 |  |
| *Cx.p.quinquefasciatus* | 1.12 | 2.9 | 0 | Transmission | 39 | 0 |  |
| *Cx.p.quinquefasciatus* | 1.13 | 2.9 | 0 | Infection | 39 | 0 |  |
| *Cx.p.quinquefasciatus* | 1.13 | 2.9 | 0 | Dissemination | 39 | 0 |  |
| *Cx.p.quinquefasciatus* | 1.13 | 2.9 | 0 | Transmission | 39 | 0 |  |
| *Cx.p.quinquefasciatus* | 1.14 | 2.9 | 0 | Infection | 39 | 0 |  |
| *Cx.p.quinquefasciatus* | 1.14 | 2.9 | 0 | Dissemination | 39 | 0 |  |
| *Cx.p.quinquefasciatus* | 1.14 | 2.9 | 0 | Transmission | 39 | 0 |  |
| *Cx.p.quinquefasciatus* | 1.15 | 2.9 | 0 | Infection | 39 | 0 |  |
| *Cx.p.quinquefasciatus* | 1.15 | 2.9 | 0 | Dissemination | 39 | 0 |  |
| *Cx.p.quinquefasciatus* | 1.15 | 2.9 | 0 | Transmission | 39 | 0 |  |
| *Cx.p.quinquefasciatus* | 1.16 | 2.9 | 0 | Infection | 39 | 0 |  |
| *Cx.p.quinquefasciatus* | 1.16 | 2.9 | 0 | Dissemination | 39 | 0 |  |
| *Cx.p.quinquefasciatus* | 1.16 | 2.9 | 0 | Transmission | 39 | 0 |  |
| *Cx.p.quinquefasciatus* | 1.17 | 2.9 | 0 | Infection | 39 | 0 |  |
| *Cx.p.quinquefasciatus* | 1.17 | 2.9 | 0 | Dissemination | 39 | 0 |  |
| *Cx.p.quinquefasciatus* | 1.17 | 2.9 | 0 | Transmission | 39 | 0 |  |
| *Cx.p.quinquefasciatus* | 1.18 | 2.9 | 1 | Infection | 39 | 170 | 2,230448921 |
| *Cx.p.quinquefasciatus* | 1.18 | 2.9 | 0 | Dissemination | 39 | 0 |  |
| *Cx.p.quinquefasciatus* | 1.18 | 2.9 | 0 | Transmission | 39 | 0 |  |
| *Cx.p.quinquefasciatus* | 1.19 | 2.9 | 0 | Infection | 39 | 0 |  |
| *Cx.p.quinquefasciatus* | 1.19 | 2.9 | 0 | Dissemination | 39 | 0 |  |
| *Cx.p.quinquefasciatus* | 1.19 | 2.9 | 0 | Transmission | 39 | 0 |  |
| *Cx.p.quinquefasciatus* | 1.20 | 2.9 | 1 | Infection | 39 | 20 | 1,301029996 |
| *Cx.p.quinquefasciatus* | 1.20 | 2.9 | 1 | Dissemination | 39 | NA |  |
| *Cx.p.quinquefasciatus* | 1.20 | 2.9 | 1 | Transmission | 39 | NA |  |
| *Cx.p.quinquefasciatus* | 1.21 | 2.9 | 0 | Infection | 39 |  |  |
| *Cx.p.quinquefasciatus* | 1.21 | 2.9 | 0 | Dissemination | 39 |  |  |
| *Cx.p.quinquefasciatus* | 1.21 | 2.9 | 0 | Transmission | 39 |  |  |
| *Cx.p.quinquefasciatus* | 1.22 | 2.9 | 1 | Infection | 39 | 100 | 2 |
| *Cx.p.quinquefasciatus* | 1.22 | 2.9 | 1 | Dissemination | 39 | NA |  |
| *Cx.p.quinquefasciatus* | 1.22 | 2.9 | 1 | Transmission | 39 | NA |  |
| *Cx.p.quinquefasciatus* | 1.23 | 2.9 | 1 | Infection | 39 | NA |  |
| *Cx.p.quinquefasciatus* | 1.23 | 2.9 | 1 | Dissemination | 39 | NA |  |
| *Cx.p.quinquefasciatus* | 1.23 | 2.9 | 1 | Transmission | 39 | NA |  |
| *Cx.p.quinquefasciatus* | 1.24 | 2.9 | 0 | Infection | 39 | 0 |  |
| *Cx.p.quinquefasciatus* | 1.24 | 2.9 | 0 | Dissemination | 39 | 0 |  |
| *Cx.p.quinquefasciatus* | 1.24 | 2.9 | 0 | Transmission | 39 | 0 |  |
| *Cx.p.quinquefasciatus* | 1.25 | 2.9 | 1 | Infection | 39 | 830 | 2,919078092 |
| *Cx.p.quinquefasciatus* | 1.25 | 2.9 | 1 | Dissemination | 39 | NA |  |
| *Cx.p.quinquefasciatus* | 1.25 | 2.9 | 1 | Transmission | 39 | NA |  |
| *Cx.p.quinquefasciatus* | 1.26 | 2.9 | 1 | Infection | 39 | 40 | 1,602059991 |
| *Cx.p.quinquefasciatus* | 1.26 | 2.9 | 0 | Dissemination | 39 | 0 |  |
| *Cx.p.quinquefasciatus* | 1.26 | 2.9 | 0 | Transmission | 39 | 0 |  |
| *Cx.p.quinquefasciatus* | 1.27 | 2.9 | 0 | Infection | 39 | 0 |  |
| *Cx.p.quinquefasciatus* | 1.27 | 2.9 | 0 | Dissemination | 39 | 0 |  |
| *Cx.p.quinquefasciatus* | 1.27 | 2.9 | 0 | Transmission | 39 | 0 |  |
| *Cx.p.quinquefasciatus* | 1.28 | 2.9 | 1 | Infection | 39 | 60 | 1,77815125 |
| *Cx.p.quinquefasciatus* | 1.28 | 2.9 | 0 | Dissemination | 39 | 0 |  |
| *Cx.p.quinquefasciatus* | 1.28 | 2.9 | 0 | Transmission | 39 | 0 |  |
| *Cx.p.quinquefasciatus* | 1.29 | 2.9 | 0 | Infection | 39 | 0 |  |
| *Cx.p.quinquefasciatus* | 1.29 | 2.9 | 0 | Dissemination | 39 | 0 |  |
| *Cx.p.quinquefasciatus* | 1.29 | 2.9 | 0 | Transmission | 39 | 0 |  |
| *Cx.p.quinquefasciatus* | 1.30 | 2.9 | 0 | Infection | 39 | 0 |  |
| *Cx.p.quinquefasciatus* | 1.30 | 2.9 | 0 | Dissemination | 39 | 0 |  |
| *Cx.p.quinquefasciatus* | 1.30 | 2.9 | 0 | Transmission | 39 | 0 |  |
| *Cx.p.quinquefasciatus* | 1.31 | 2.9 | 0 | Infection | 39 | 0 |  |
| *Cx.p.quinquefasciatus* | 1.31 | 2.9 | 0 | Dissemination | 39 | 0 |  |
| *Cx.p.quinquefasciatus* | 1.31 | 2.9 | 0 | Transmission | 39 | 0 |  |
| *Cx.p.quinquefasciatus* | 1.32 | 2.9 | 0 | Infection | 39 | 0 |  |
| *Cx.p.quinquefasciatus* | 1.32 | 2.9 | 0 | Dissemination | 39 | 0 |  |
| *Cx.p.quinquefasciatus* | 1.32 | 2.9 | 0 | Transmission | 39 | 0 |  |
| *Cx.p.quinquefasciatus* | 1.33 | 2.9 | 1 | Infection | 39 | NA |  |
| *Cx.p.quinquefasciatus* | 1.33 | 2.9 | 1 | Dissemination | 39 | 720 | 2,857332496 |
| *Cx.p.quinquefasciatus* | 1.33 | 2.9 | 1 | Transmission | 39 | NA |  |
| *Cx.p.quinquefasciatus* | 1.34 | 2.9 | 1 | Infection | 39 | NA |  |
| *Cx.p.quinquefasciatus* | 1.34 | 2.9 | 0 | Dissemination | 39 | 0 |  |
| *Cx.p.quinquefasciatus* | 1.34 | 2.9 | 0 | Transmission | 39 | 0 |  |
| *Cx.p.quinquefasciatus* | 1.35 | 2.9 | 1 | Infection | 39 | 140 | 2,146128036 |
| *Cx.p.quinquefasciatus* | 1.35 | 2.9 | 0 | Dissemination | 39 | 0 |  |
| *Cx.p.quinquefasciatus* | 1.35 | 2.9 | 0 | Transmission | 39 | 0 |  |
| *Cx.p.quinquefasciatus* | 1.36 | 2.9 | 0 | Infection | 39 | 0 |  |
| *Cx.p.quinquefasciatus* | 1.36 | 2.9 | 0 | Dissemination | 39 | 0 |  |
| *Cx.p.quinquefasciatus* | 1.36 | 2.9 | 0 | Transmission | 39 | 0 |  |
| *Cx.p.quinquefasciatus* | 1.37 | 2.9 | 0 | Infection | 39 | 0 |  |
| *Cx.p.quinquefasciatus* | 1.37 | 2.9 | 0 | Dissemination | 39 | 0 |  |
| *Cx.p.quinquefasciatus* | 1.37 | 2.9 | 0 | Transmission | 39 | 0 |  |
| *Cx.p.quinquefasciatus* | 1.38 | 2.9 | 0 | Infection | 39 | 0 |  |
| *Cx.p.quinquefasciatus* | 1.38 | 2.9 | 0 | Dissemination | 39 | 0 |  |
| *Cx.p.quinquefasciatus* | 1.38 | 2.9 | 0 | Transmission | 39 | 0 |  |
| *Cx.p.quinquefasciatus* | 1.39 | 2.9 | 0 | Infection | 39 | 0 |  |
| *Cx.p.quinquefasciatus* | 1.39 | 2.9 | 0 | Dissemination | 39 | 0 |  |
| *Cx.p.quinquefasciatus* | 1.39 | 2.9 | 0 | Transmission | 39 | 0 |  |

ID: Individuals identification. Viremia ( $\log_{10}$ PFU/ml). Results: 0 negative, 1 positive. Infection rate (number of positive mosquito abdomens/total number of mosquitoes fed), dissemination rate (number of mosquitoes with positive legs/total number of mosquitoes fed) and transmission rate (number of mosquitoes with positive saliva/total number). N: Total number of mosquitoes assayed. SLEV titers in abdomen, legs and saliva of each mosquito species in PFU/ml and $\log_{10}$ PFU/ml. NA: No data.

